# Micropropagated walnut dependency on phosphate fertilization and arbuscular mycorrhiza for growth, nutrition and quality differ between rootstocks both after acclimatization and *post*-acclimatization

**DOI:** 10.1101/2022.11.24.517850

**Authors:** Emma Mortier, Samuel Jacquiod, Laurent Jouve, Fabrice Martin-Laurent, Ghislaine Recorbet, Olivier Lamotte

## Abstract

The English walnut (*Juglans regia* L.) is the main species cultivated for the production of edible nuts. *In vitro* micropropagation of walnut explants, necessary for mass propagation of high-quality walnut rootstocks, needs an *ex vitro* acclimatization phase followed by a *post*-acclimatization growth in greenhouse when plantlets become photoautotrophic. However, poor survival and slow growth rates are common difficulties encountered in nurseries when establishing micropropagated walnut saplings. As many other fruit and nut bearing trees, walnut exhibits a high dependency on symbiotic soil-borne arbuscular mycorrhizal fungi for better soil nutrient acquisition and development due to a coarse root architecture that notably limits soil inorganic phosphate (Pi) uptake. In the context of rootstock production, we investigated the establishment of seven walnut rootstocks of economic interest (RG2, RG6, R17, RX1, VLACH, VX211, and WIP3) inoculated or not with *Rhizophagus irregularis* under two contrasting Pi fertilization regimes. We demonstrated that inoculation with *R. irregularis* decreases micropropagated walnut rootstock dependency on Pi fertilization both at the acclimatization and *post*-acclimatization stages, together with improving quality, sapling biomass production and nutrition of walnut rootstocks. We also showed that these benefits are rootstock-dependent, indicating that walnut mycorrhizal dependency for Pi nutrition varies between cultivars.

## Introduction

Walnut is the common name given to twenty-five species of deciduous trees belonging to the genus *Juglans* (order Fagales, family Juglandaceae), which are distributed across the temperate and subtropical regions of the northern hemisphere [1]. All species produce nuts, but *J. regia*, the English or Persian walnut, is the main species widely cultivated for nut production. Obtaining walnut trees is commonly carried out by grafting selected varieties of English walnut scions onto seedling rootstocks selected to provide the best anchorage, vigour, and tolerance of pathogens [2]. As such, the primary commercial importance of hybrid rootstocks ‘Paradox’ (*J. hindsii* x *J. regia*) is to provide tolerance of most unfavourable soil conditions as well as being relatively resistant to *Phytophthora* spp. and nematodes [3]. Because walnut is difficult to propagate clonally from cuttings, *in vitro* plant tissue culture has increased over the past decades and plays a key role in mass propagation of high-quality walnut cultivars and rootstocks [4]. The first *in vitro* propagation procedures for walnut trees were developed more than thirty years ago for the hybrid rootstock ‘Paradox’ [5]. Advances in micropropagation and breeding technologies have further enabled the exploration of diverse *Juglans* species and hybrids for the improvement of walnut rootstocks [6]. A major hurdle in adopting clonal propagation for walnuts is that explants are much more difficult to root than rootstocks of other fruit and nut crops [7], and further undergo restricted inorganic phosphate (Pi) nutrition that limits biomass production [8]. In addition, saturated atmosphere and low light intensity on artificial culture media cause developmental distortions [7], including a root system devoid of root hairs, which limits nutrient and water uptake, poor cuticle wax development, a low chlorophyll content, non-functional stomata and reduced photosynthetic efficiency [9]. Consequently, when transplanted *ex vitro*, micropropagated plantlets suffer from severe environmental stresses and substantial losses may occur during acclimatization to the *ex vitro* environment. Walnut explants therefore need an acclimatization phase to repair the *in vitro* induced abnormalities and further require a *post*-acclimatization growth in greenhouse conditions when plantlets become photoautotrophic [9–10]. In this regard, pre-planting quality assessment is the keystone event in tree nursery production, because low nutrient availability resulting from poor root soil contact, low water porosity of suberized roots, and mechanical root damage are responsible for low survival of transplanted seedlings [11–12]. According to Fonseca et al. [13], variables used for evaluating seedling quality should not be studied separately, that way avoiding the risk of selecting higher and yet weaker seedlings while discarding smaller, sturdier ones. Therefore, the Dickson quality index [14] is considered as a reliable indicator of seedling quality because its calculation computes robustness and biomass distribution [15].

In their natural environment, plants are colonized by beneficial microorganisms that improve plant growth and health under abiotic and biotic stresses [16]. The majority of woody and fruit trees, including walnut, depend on symbiotic soil-borne arbuscular mycorrhizal (AM) fungi for normal growth and development due to the morphology of the root system, typically with coarse roots and relatively few roots per plant [17]. AM fungi, which belong to an ancient lineage of obligate biotrophs in the sub-phylum Glomeromycotina [18], colonize plant roots to obtain plant-derived carbon in the form of sugars and lipids for their growth and reproduction [19]. In return, AM fungi provide soil mineral nutrients, mainly Pi, to the host, which are acquired through the fungal extra-radical mycelium that reaches soil volumes inaccessible to plant roots [19–20]. The stimulating effects of AM fungi on plant growth and physiology include a higher branching of the root system, an improved nutritional state due to an increased Pi supply, which in turn leads to increased photosynthetic rates and improved plant growth [16]. Such characteristics have led to considerable interest in evaluating the significance of inoculating micropropagated plantlets with selected AM fungi to contribute to their biohardening during the acclimatization and *post*-acclimatization stages [21]. The beneficial effect of AM symbiosis on walnut quality seedling and field performance has been reported for the eastern black walnut *J. nigra* for which mycorrhization enhances transplant success by increasing the number of lateral roots and plant P uptake [22–23]. More recently, application of AM fungi on 1-year-old *J. regia* seedlings was found to improve plant growth and nutrition under drought stress [24]. Likewise, Huang et al. [25] demonstrated that mycorrhization of germinated *J. regia* seedlings has positive effects on plant growth, root morphology, leaf gas exchange, and root nutrient acquisition after three months. However, the later studies were conducted on saplings grown from seedlings, which may be of limited relevance for micropropagated walnut plantlets because most of roots found during the first stage of acclimatization are *in vitro*-formed [6, 25].

To the best of our knowledge, only a few number of studies have addressed the impact of AM fungi on the acclimatization and post-acclimatization of micropropagated *Juglans* spp. Peixe et al. [10] showed that inoculation with *Glomus* spp. did not improve the *ex vitro* survival of micropropagated *J. regia* x *J. hindsii* rootstocks, while Dolcet-Sanjuan et al. [26] reported that inoculation of micropropagated *J. regia* with the AM fungus *Glomus mosseae* or *Rhizophagus irregularis* significantly improved plant survival when transferred to nursery. By contrast, no data are available regarding the mycorrhizal behaviour of micropropagated walnut explants upon their pre-inoculation with an AM fungus on seedling quality, mineral nutrition, and growth responses, as referred to as mycorrhizal responsiveness and mycorrhizal dependency. Mycorrhizal responsiveness relates directly to host-growth rate and internal Pi demand, while mycorrhiza dependency is more related to the nutrient absorption ability of a non-colonized plant than to its nutrient requirement [27–28]. In this context, it is noteworthy that these mycorrhizal growth traits are determined by soil phosphate availability, but is also an intrinsic property of a plant species or genotype largely controlled by plant growth rate and the root system architecture [27]. Mycorrhizal responses are for example negatively correlated with root morphological characteristics such as root length, root hair length, and density of root hairs [29]. Because the rooting ability of micropropagated walnut plantlets is genotype-dependent [6, 26], further research is needed to assess the potential of the commercially available AM fungus *R. irregularis* to improve the acclimatization and *post*-acclimatization of micropropagated walnut explants as related to the impact of Pi supply and rootstock cultivaron their mycorrhizal seedling quality, nutrition, and growth responses. To address these questions, we set up two experiments in which the mycorrhizal behaviour of seven walnut rootstocks were recorded upon two contrasting Pi fertilization regimes during the acclimatization and *post*-acclimatization stages.

## Results

### Experiment 1. Nutrition, growth and quality of micropropagated walnut rootstocks after acclimatization as dependent on Pi fertilization and pre-inoculation with *R. irregularis*

Mycorrhizal root colonization parameters were recorded after two months of acclimatization in plants pre-inoculated (AM) or not (NM) with the AM fungus *R. irregularis* supplied with two contrasting levels of Pi fertilization **(Figure S1)**. As expected, no fungal structures were observed in NM walnut roots. In AM plants, mycorrhization developed to a similar extent in the two fertilization regimes in the rootstocks RG2, RG6, RG17, RX1 VX211 and WIP3: mycorrhization parameters F, M and A% ranged from 30 to 100%, 1.3 to 20%, and 0.9 to 15%, respectively (**Figure 1**). By contrast, in the rootstock VLACH, F, M and A% were significantly lower in the highest Pi treatment: when compared to the M and A% that reached 21 and 18% in the P/10 condition, less than 2% of VLACH root length was colonized by intraradical hyphae, and only 1% root length formed arbuscules in the PTOT condition. These results indicate that both rootstock and Pi supply have an influence on the mycorrhizal colonization after the acclimatization stage.

**Figure 1.**
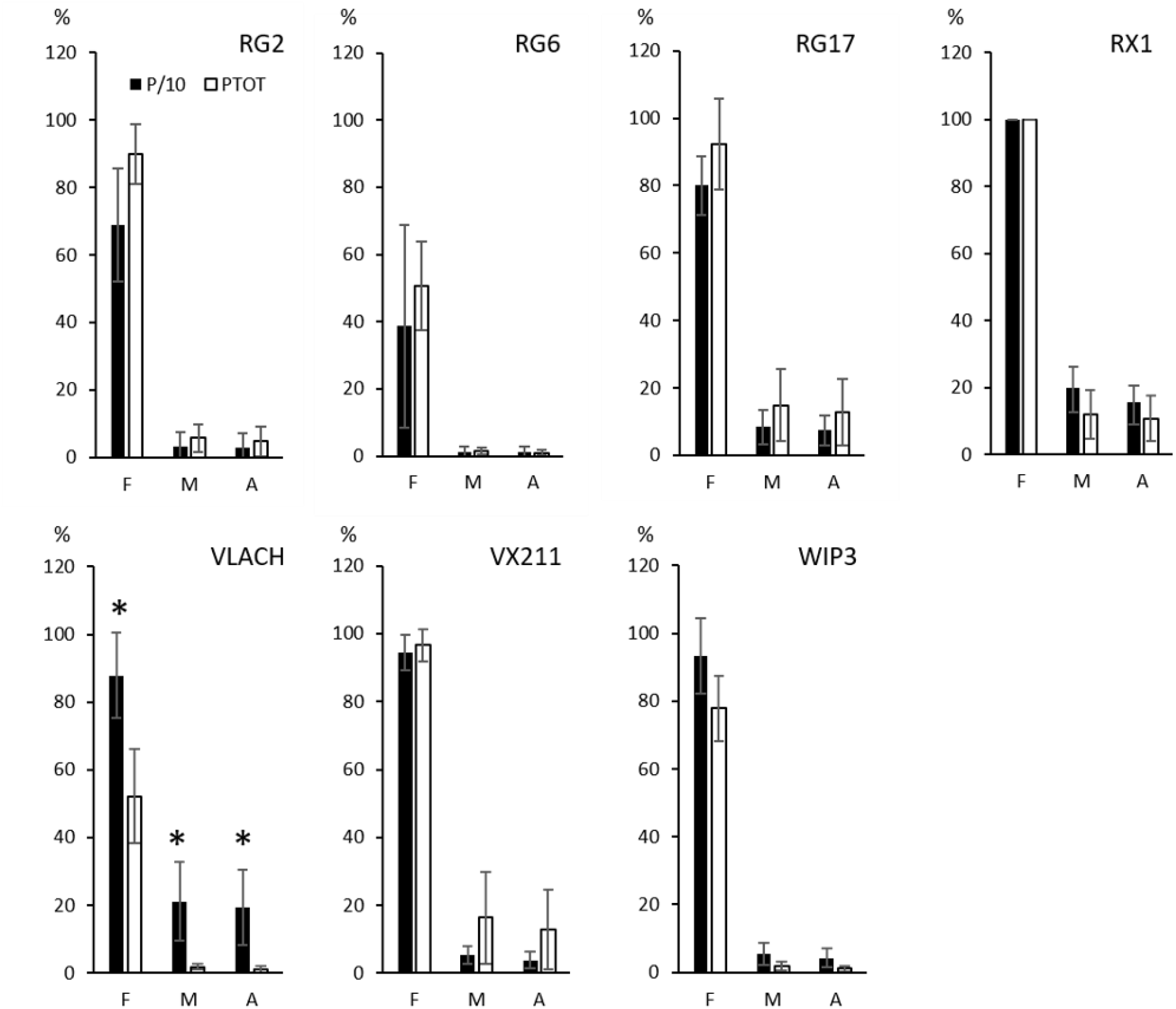
Mycorrhizal root colonization after two months of acclimatization of walnut rootstocks pre-inoculated with the AM fungus *R. irregularis* under two contrasting Pi fertilization regimes. Mean data (±SE) of height to 15 replicates per mycorrhizal treatment are presented in the P/10 (dark bars) and PTOT (white bars) fertilization regimes. Asterisks indicate significant (*p* < 0.05) differences between the two Pi fertilization regimes. F%: mean external frequency of mycorrhization of root fragments, M%: intensity of root cortex colonization, A%: mean arbuscular abundance.

To investigate the effects of pre-inoculation with *R. irregularis* on walnut rootstocks at the end of the acclimatization stage, growth, quality and nutritional parameters were first compared between AM and NM plants under the two contrasting fertilization regimes. The data displayed in **Table S1** show that for the rootstock RG2, leaf Pi concentrations were significantly lower in the P/10 treatment than in the PTOT condition in NM and AM plants. Irrespective of the Pi supply, mycorrhization did not affect the development and the nutrition of RG2, but the root dry weight (DW) was significantly higher in AM-PTOT plants than in AM-P/10 plants. Pre-inoculation of RG6 had no impact in the P/10 condition, but led to a significant increase in root DW, and rootstock quality, as measured by the Dickson quality index (DQI) in the PTOT treatment. In the two Pi fertilization regimes, root DW and DQI were higher in mycorrhizal than in non-mycorrhizal RG17 rootstocks. In parallel, AM RG17 and RX1 walnuts showed a significantly lower shoot-to-root (S/R) DW ratio than non-inoculated plants whatever the Pi supply. Under the PTOT condition, pre-inoculation of VX211 resulted in a significant increase in leaf Pi concentration without affecting the rootstock morphological parameters. In the P/10 treatment, pre-inoculation of VX211 resulted in a higher the Pi concentration and a lower S/R DW ratio relative to NM plants. Irrespective of mycorrhization, significantly lower primary root lengths and root surfaces, together with higher shoot DW and S/R DW ratios were recorded in the P/10 condition relative to the PTOT treatment. In the two fertilization regimes, mycorrhization of WIP3 led to a significant increase in leaf Pi concentration without significantly affecting morphological parameters. In both AM and NM WIP3 plants, height and root surface were lower under the P/10 condition than under the PTOT fertilization.

With regard to the rootstock VLACH, pre-inoculation resulted in higher leaf Pi concentrations relative to NM plants in the two fertilization regimes with different outcomes on morphological parameters (**Figure 2A**). In the PTOT treatment, the primary root length was lower in AM than in NM rootstocks (**Figure 2G**). In the P/10 condition, the collar, DQI, root, shoot and total DWs were significantly higher in AM than NM plants (**Figure 2H, I, B, C, D**). Relative to the NM-PTOT treatment, the NM-P/10 condition resulted in a significant decrease in primary root length, height, root DW, shoot DW, total DW and explant quality (**Figure 2G, F, B, C, D, I**), indicating that the lower Pi availability in the P/10 treatment limited the development, quality and nutrition of VLACH. Noticeably, **Figure 2** showed that AM symbiosis compensated the reduced Pi availability in the P/10 treatment for VLACH, as inferred from the similar values recorded between the AM-P/10 and NM-PTOT conditions for parameters such as plant height, root DW, shoot DW, total DW, and quality index (**Figure 2F, B, C, D, I**).

**Figure 2.**
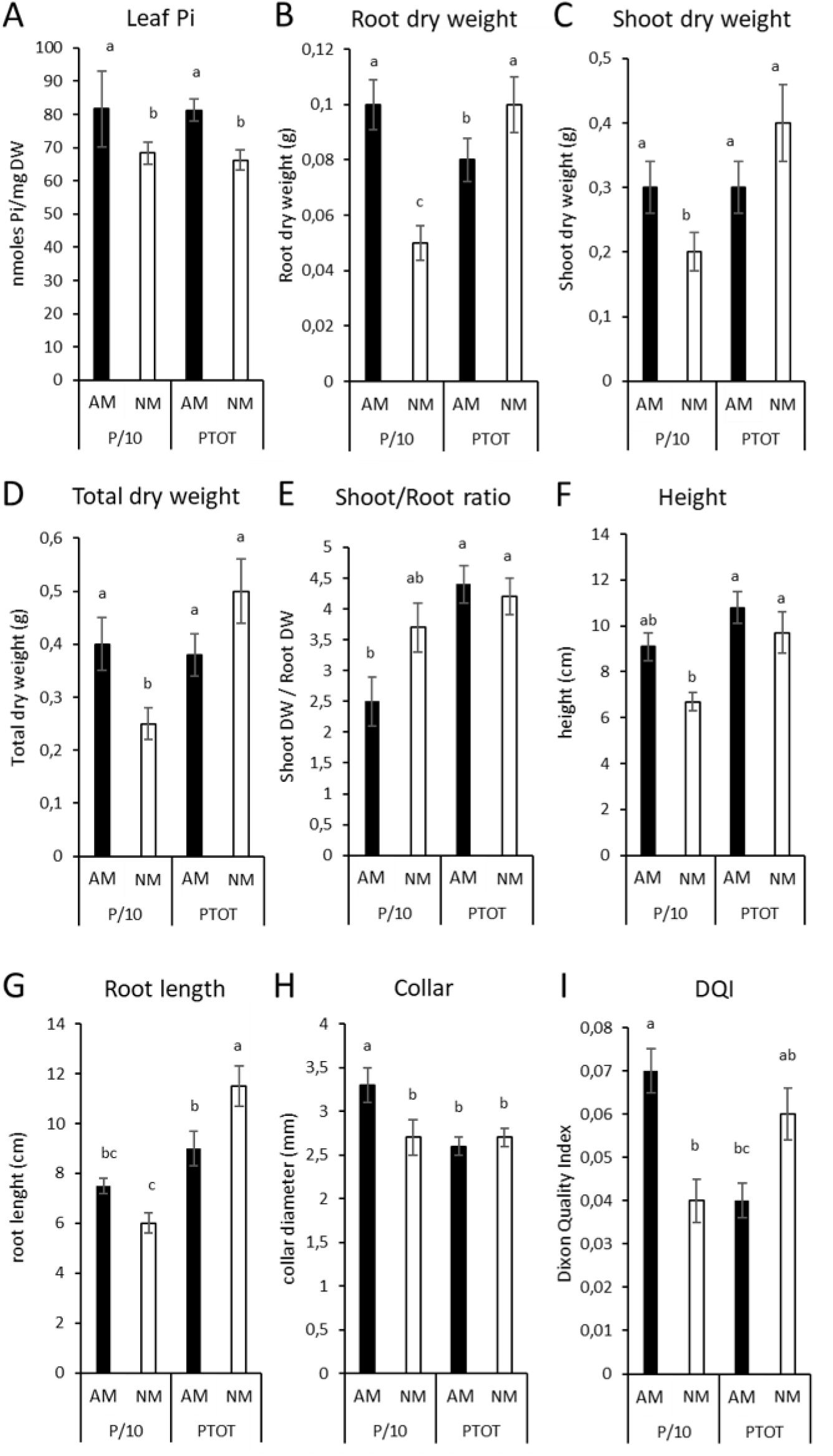
Growth, nutrition and quality responses of VLACH walnut rootstock to pre-inoculation with *R. irregularis* after two months of acclimatization under two different Pi supplies. Mean data (±SE) of height to 15 replicates per treatment are presented in the P/10 (dark bars) and PTOT (white bars) fertilization regimes. Different letters indicate statistical differences between groups (*p* < 0.05) calculated as described in the material and methods section. AM: mycorrhizal; DW: dry weight; NM: non-mycorrhizal.

To compare the impact of the differential Pi supply on the walnut overall behaviour in response to pre-inoculation with *R. irregularis* as dependent on the rootstock cultivar, we calculated in the two Pi fertilization regimes the mycorrhizal dependency (MD%), the mycorrhizal growth response (MGR%), the mycorrhizal shoot phosphate response (MPRS%) and the mycorrhizal quality response (MQR%). **Table S2** showed that rootstock responses to pre-inoculation were variable depending on the rootstock and the Pi supply, ranging from negative, neutral, positive to highly positive, as indicated by blue, grey, pink and dark pink-shaded values, respectively. These were further explored through a principal component analysis (PCA) to assess the relationships between the different parameters. As displayed in **Figure 3A**, the first principal component (PC1), which explained 74.2% of the variation in the data, captured most of the variance in terms of walnut mycorrhizal responses. Noticeably, MD, MGR, MPRS, MQR% and P/10 fertilization were positively correlated to each other and negatively correlated to the highest Pi fertilization regime on this first axis. As shown in **Figure 3B**, in the lowest Pi supply (P/10: blue circles), the mycorrhizal behaviour of RG2, VLACH, VX211 and WIP3 clustered to the positive side of PC1, opposite to what observed upon the highest Pi fertilization regime (PTOT: green circles). On the contrary, the mycorrhizal responses of RG6 and RG17 increased in the PTOT condition relative to the P/10 treatment. Taken together, these results show that after acclimatization, shoot phosphate nutrition, growth and quality responses to pre-inoculation with *R. irregularis* depend on the rootstock and the level of Pi supply.

**Figure 3.**
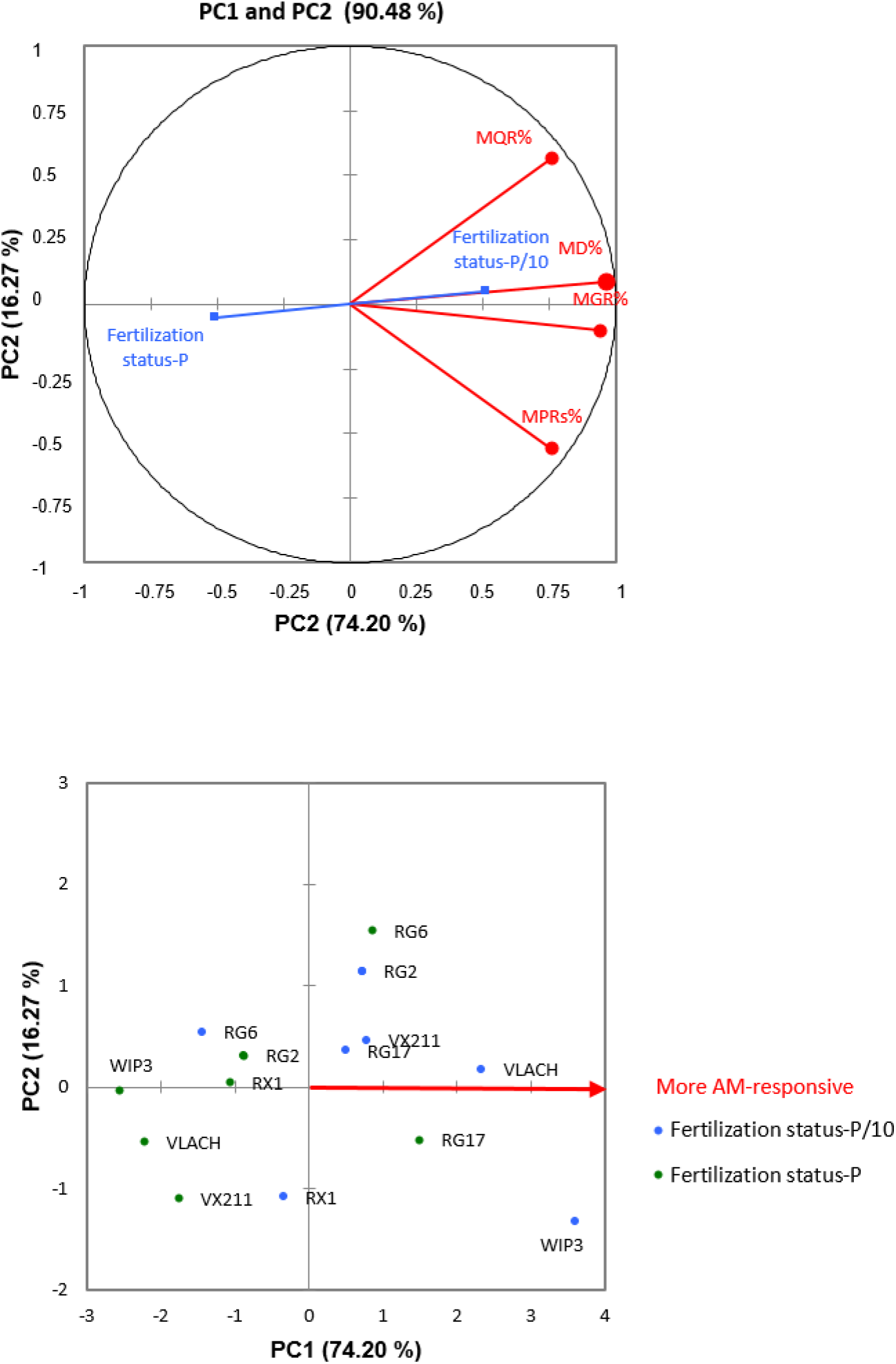
Principal Component Analysis (PCA) showing the relationships between walnut rootstock responses to pre-inoculation with *R. irregularis* after two months of acclimatization under two different Pi supplies. (A) Circle of correlations showing that the positive side of axis PC1 is correlated to walnut responses to *R. irregularis* in the lowest Pi supply. Red and blue colours refer to active and supplementary variables, respectively. (B) Responses to *R. irregularis* inoculation of walnut rootstocks under contrasting Pi fertilization regimes. Blue and green colours refer to P/10 and PTOT supply, respectively. Values correspond to the mean of height to 15 replicates per mycorrhizal and non-mycorrhizal plants for each Pi supply. MD%, mycorrhizal dependency; MGR%, mycorrhizal growth response, MPRs%, mycorrhizal shoot phosphate response, MQR%, mycorrhizal quality response.

### Experiment 2. Nutrition, growth and quality of micropropagated walnut rootstocks after *post*-acclimatization as dependent on Pi fertilization and pre-inoculation with *R. irregularis*

After acclimatization of the non-mycorrhizal rootstocks, walnut plants were either pre-inoculated or not with *R. irregularis* and further *post*-acclimatized in growth chamber conditions using two different Pi fertilization regimes (**Figure S1**). Fungal structures recorded after two months of *post*-acclimatization were not observed inside and outside the roots of NM rootstocks. The results displayed in **Figure 4** show that fungal structures developed to a similar extent in the two Pi supplies in the rootstocks RG2, RG6, RG17, RX1, VLACH and WIP3. Depending on the rootstock analysed, F, M, and A% ranged from 60 to 99%, 1.4 to 21%, and 0.7 to 13%, respectively. By contrast, in the rootstock VX211, the three parameters were significantly lower in the highest Pi treatment. Compared to the M and A% that reached 9 and 8% in the P/10 condition, less than 4% of VX211 root length was colonized by intraradical hyphae and only 1.6% root length formed arbuscules in the PTOT condition. These results indicate that both rootstock and Pi supply have an influence on mycorrhizal colonization after the *post*-acclimatization stage.

**Figure 4.**
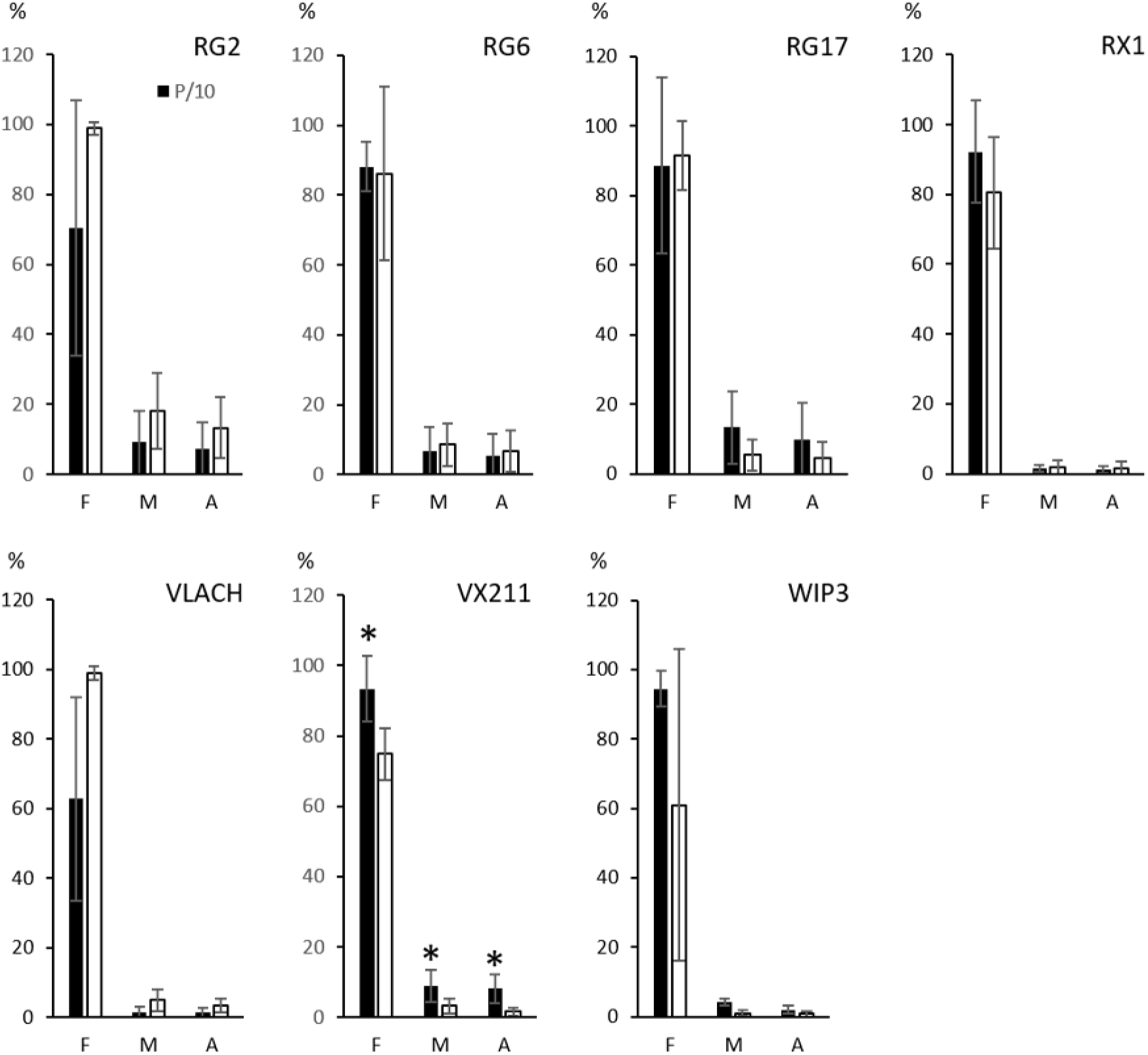
Mycorrhizal root colonization after two months of *post*-acclimatization of walnut rootstocks pre-inoculated with the AM fungus *R. irregularis* under two contrasting Pi fertilization regimes. Mean data (±SE) of height to 15 replicates per mycorrhizal treatment are presented in the P/10 (dark bars) and PTOT (white bars) fertilization regimes. Asterisks indicate significant (*p* < 0.05) differences between the two Pi fertilization regimes. F%: mean external frequency of mycorrhization of root fragments, M%: intensity of root cortex colonization, A%: mean arbuscular abundance.

Growth, quality and nutritional parameters were further compared between AM and NM plants under the two contrasting Pi regimes in order to assess the effects of pre-inoculation with *R. irregularis* on walnut rootstocks after the *pos*t-acclimatization stage. **Table S3** showed that under the two Pi supplies, AM RG2 rootstocks, relative to the respective NM controls, displayed a higher leaf Pi concentration, together with a lower primary root length. In both AM and NM RG2, root DW, total DW, DQI were significantly lower, and the S/R DW ratio significantly higher in the P/10 condition than in the PTOT condition. Under the two fertilization regimes, pre-inoculation of RG6 did not significantly impact the development and nutrition of the rootstocks relative to NM saplings. In both AM and NM RG6, S/R DW ratios were significantly higher in the P/10 condition than in the PTOT treatment. Under the two Pi supplies, AM RG17 rootstocks, relative to NM controls, displayed an increased height and root Pi concentration. In both AM and NM RG17 rootstocks, root and leaf Pi concentrations were inferior in the P/10 condition relative to the PTOT fertilization. With regard to RX1, whatever the fertilization, pre-inoculation did not significantly impact the nutritional parameters relative to NM saplings, but significantly increased rootstock quality. AM RX1 displayed a higher total DW in the P/10 treatment and a higher shoot DW in the PTOT condition. Irrespective of the mycorrhizal status, a significantly higher root surface was recorded in the P/10 condition relative to the PTOT condition. Under the two fertilization regimes, pre-inoculation of VLACH resulted in a significant increase in the root Pi concentration without impacting morphological parameters. In both AM and NM VLACH saplings, root DW and DQI were higher in the lowest fertilization. Under the lower Pi supply (P/10) relative to NM controls, AM VX211 displayed a significantly increase in root and leaf Pi concentrations without any significant effect on morphological parameters. In this rootstock, irrespective of the mycorrhizal status, root surfaces, shoot and root Pi concentrations were significantly lower under the P/10 fertilization than in the PTOT treatment, while S/R DW ratios were higher. Relative to NM rootstocks, pre-inoculation of WIP3 with *R. irregularis* under the P/10 condition resulted in increased height, branch number, root Pi concentration together with a lower root surface (**Figure 5E, F, A, C**). In the PTOT treatment, AM WIP3 displayed a higher height and branch number than NM rootstocks (**Figure 5E, F**). Irrespective of the mycorrhizal status, root surfaces were higher, while root Pi concentrations were lower in the P/10 than in the PTOT condition (**Figure 5C, A**). Noticeably, the total DW of AM WIP3 was higher in the P/10 than in the PTOT fertilization (**Figure 5G**).

**Figure 5.**
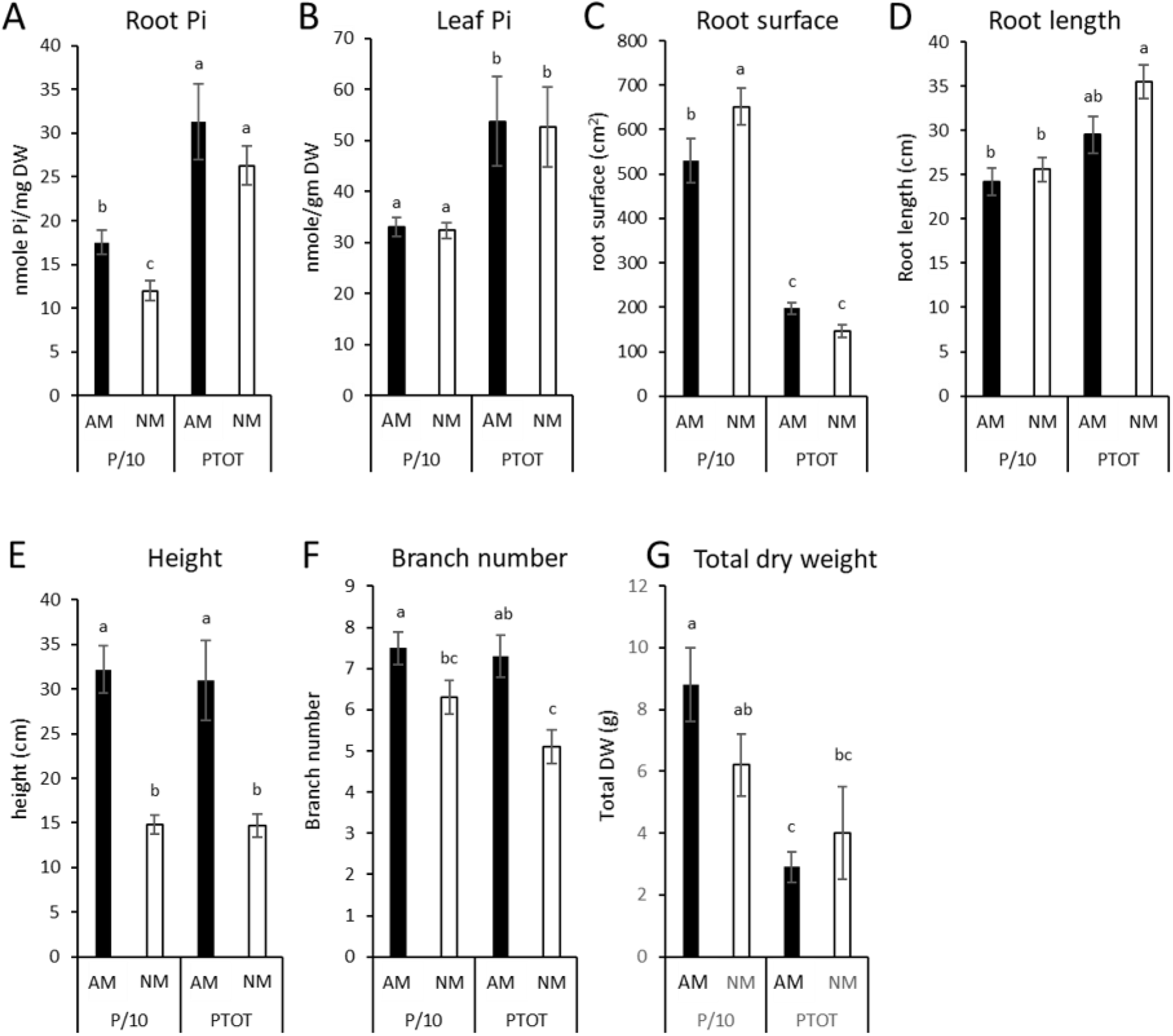
Growth, nutrition and quality responses of WIP3 walnut rootstock to pre-inoculation with *R. irregularis* after two months of *post*-acclimatization under two different Pi supplies. Mean data (±SE) of height to 15 replicates per treatment are presented in the P/10 (dark bars) and PTOT (white bars) fertilization regimes. Different letters indicate statistical differences between groups (*p* < 0.05) calculated as described in the material and methods section. AM: mycorrhizal; DW: dry weight; NM: non-mycorrhizal.

The comparison of growth, quality and nutritional parameters in the non-inoculated walnut plants between the two Pi supplies, showed that the NM-P/10 condition relative to the NM-PTOT treatment, resulted in a significant decrease in plant primary root length (WIP3), collar diameter (RG2), root DW (RG2), total dry biomass (RG2, RX1), DQI (RG2), root surface (VX211), root Pi concentration (RG17, VX211,WIP3), leaf Pi concentration (RG17, VLACH, VX211) (**Table S3**). These results indicate that the reduced Pi availability in the P/10 condition limited the development, quality, and/or the nutrition of the corresponding rootstocks. The analysis of these parameters between the NM-PTOT, NM-P/10 and AM-treatments (**Table S3**), underlined that pre-inoculation with *R*. *irregularis* after the acclimatization stage, either totally (RG17 root Pi concentration; RX1 total DW, VLACH leaf Pi concentration), or partially (VX211 root and leaf Pi concentrations, WIP3 root Pi concentration) alleviated Pi deficiency. For the other parameters, pre-inoculation with *R. irregularis* did not significantly rescue low Pi availability, especially in the rootstocks RG2 and RG6 (**Table S3**). Taken together, these results demonstrate that establishment of AM symbiosis alleviates low Pi availability by significantly restoring or improving the nutritional status in VLACH, VX211 and WIP3 rootstocks.

To further compare the impact of the Pi supply on the response of different micropropagated rootstocks to pre-inoculation with *R. irregularis* at the *post*-acclimatization stage, we analysed in the two Pi fertilization regimes the relationship between MD, MGR MQR, mycorrhizal shoot phosphate (MPRS) and root phosphate (MPRR) responses. After the *post*-acclimatization stage, walnut responses to pre-inoculation, as dependent on the rootstock and the Pi supply, ranged from negative, neutral, positive to highly positive, as indicated by blue, grey, pink and dark pink-shaded values, respectively (**Table S4**). After a PCA to assess the relationships between the different parameters, **Figure 6A** showed that PC1 that explained 48.9% of the variation in the data, captured mycorrhizal responses with regard to MD, MGR, and MQR%. Noticeably, these parameters were positively correlated to each other and to the lowest Pi fertilization regime on this first axis. **Figure 6B** indicated that in the lowest Pi supply (P/10: blue circles), the mycorrhizal behaviour of VLACH, VX211 and WIP3 clustered to the positive side of PC1, opposite to what observed upon the highest Pi fertilization regime (P, green circles). Irrespective of the Pi fertilization regime, MGR, MD and MR% were negative for RG2, while positive for RX1. Mycorrhizal root phosphate response (MPRR) loaded positively on PC2 that explained 30% of the variation (**Figure 6A)**. The latter axis was essentially driven by the mycorrhizal shoot phosphate response of RX1 that clustered to the positive side of PC2 in the lowest Pi supply, opposite to what observed upon the highest Pi fertilization regime (**Figure 6B**). A similar trend was also observed for the rootstock WIP3. On PC1 and PC2, the overall mycorrhizal behaviour of RX1, VLACH and WIP3 were superior in the P/10 treatment than in the PTOT condition. Taken together, these results show that after *post*-acclimatization, both rootstock and Pi supply influenced the response of walnut rootstock to *R. irregularis*.

**Figure 6.**
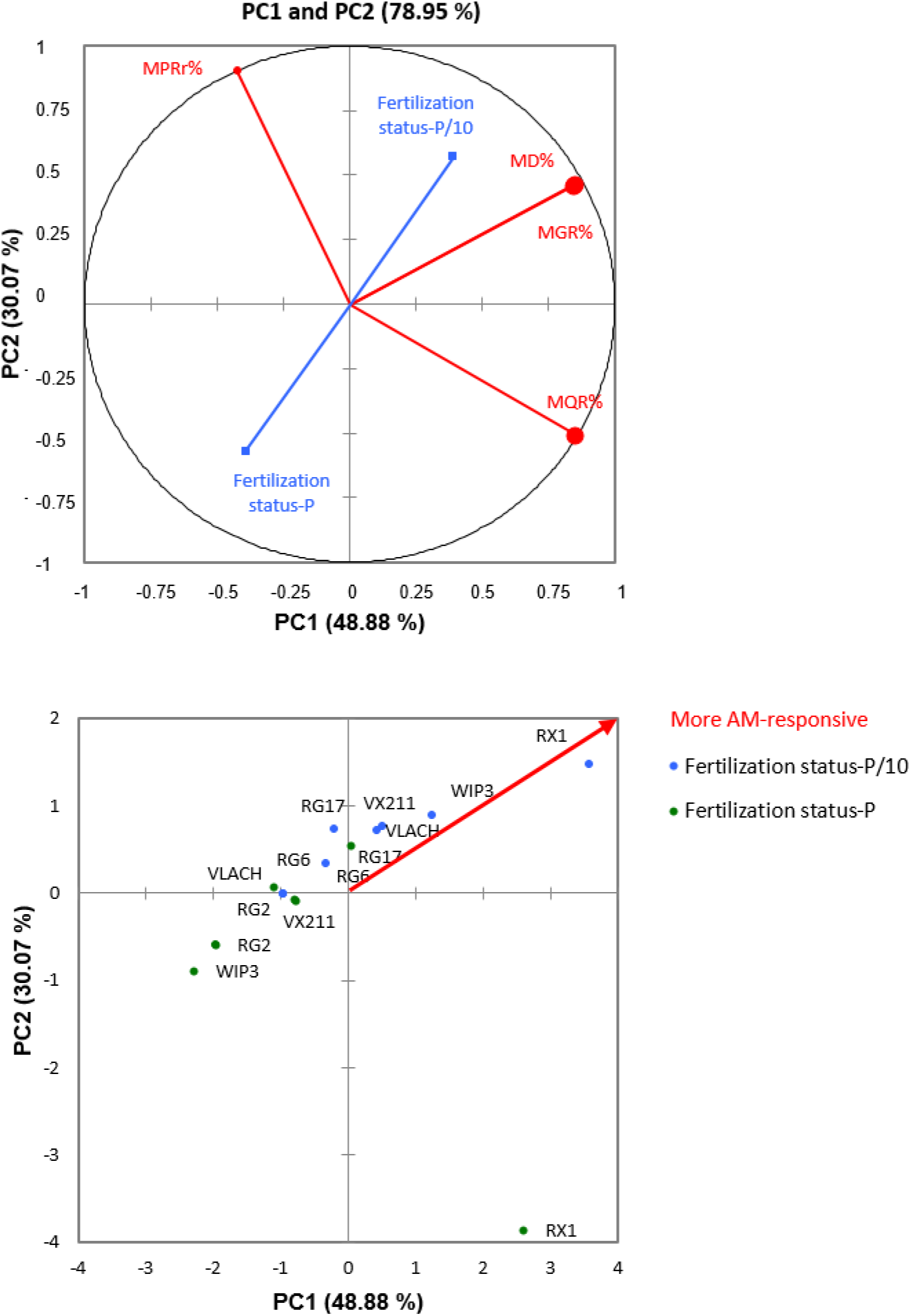
Principal Component Analysis (PCA) of the relationships between walnut rootstock responses to pre-inoculation with *R. irregularis* after two months of *post*-acclimatization under two different Pi supplies. (A) Circle of correlations showing that the positive side of axis PC1 is correlated to walnut responses to *R. irregularis* in the lowest Pi supply, with regard to MD, MGR and MQR values. On the contrary, root phosphate response (MPRR) loaded positively on PC2. Red and blue colours refer to active and supplementary variables, respectively. (B) Responses to *R. irregularis* inoculation of walnut rootstocks under contrasting Pi fertilization regimes. Blue and green colours refer to P/10 and PTOT supply, respectively. Values correspond to the mean of height to 15 replicates per mycorrhizal and non-mycorrhizal plants for each Pi supply. MD%, mycorrhizal dependency; MGR%, mycorrhizal growth response, MPRr%, mycorrhizal root phosphate response, MQR%, mycorrhizal quality response.

## 4. Discussion

Production of high quality, marketable walnut saplings is commonly carried out by grafting selected varieties of English walnut scions onto rootstocks obtained by micropropagation, a process that needs improvements owing to substantial rootstock losses after walnut *ex vitro* establishment [3, 6]. In spite of occasional reports on the impact of biohardening with AM fungi [10, 26], little information is available regarding the growth, nutritional and sapling quality responses of walnut rootstocks pre-inoculated with *R. irregularis* after acclimatization and *post*-acclimatization. This study was therefore designed to assess the extent to which pre-inoculation with the commercially available AM fungus *R. irregularis* DAOM197198 may improve the acclimatization and *post*-acclimatization of micropropagated walnut explants as related to the impact of Pi supply and rootstock cultivar.

Root mycorrhizal parameters were first recorded to assess the degree of interaction between AM fungi and the host plant [30]. Consistent with previous values obtained in *Juglans* spp. [22], external frequency of mycorrhization of root fragments (F%) and arbuscule abundance (A%), which represent root-intrinsic fungal structures involved in nutrient exchange, ranged from 30 to 100%, and 0.9 to 15%, respectively. Although less than 3% of A% were detected in the rootstock WIP3 after *post*-acclimatization under the P/10 fertilization regime, the benefit of mycorrhization relative to NM plants, could be demonstrated in terms of increased height, root Pi concentration, and higher biomass production. These results agree with earlier studies reporting improved biomass yield and/or mineral nutrition due to mycorrhization of micropropagated plantlets of other fruit trees such as *Annona cherimola, Malus pumila, Prunus avium, Prunus cerasifera* and *Vitis vinifera* [31–33].

A negative impact of Pi supply on endomycorrhizal development was observed in the rootstocks VLACH and VX211, as dependent on their development stage. Namely, the highest Pi fertilization regime significantly depressed the mycorrhizal colonization of VLACH after acclimatization but not after *post*-acclimatization, opposite to what observed for VX211. According to Siqueira and Saggin-Júnior [28], host nutrient requirements and their fulfilment in the absence of AM fungi are the major factors determining Pi effect on root colonization of woody plants. Consistently, after acclimatization, non-mycorrhizal VLACH contrary to NM VX211, underwent biomass limitation under the P/10 fertilization regime (height, root DW, total DW), which were mostly relieved by mycorrhization. On the opposite, after *post*-acclimatization, non-mycorrhizal VX211 contrary to NM VLACH, displayed nutritional deficiencies under the P/10 fertilization regime (root and leaf Pi concentrations), which were partially compensated by mycorrhization. It has long been known that in response to high Pi levels in soil plants suppress the development of AM symbiosis [34], a phenomenon referred to as Pi inhibition of mycorrhizal colonization [35]. The negative effect of high soil Pi levels on AM formation has been ascribed to high Pi concentrations in the roots [36], leading to a reduced carbon allocation to the AM fungus after several weeks of symbiotic functioning [37]. Although this hypothesis could not be tested after acclimatization due to tiny amounts of root materials, one could observe that inoculated VX211 walnuts after the *post*-acclimatization stage displayed the highest mean root Pi concentration (45.9 nmole Pi/mg DW) relative to the values recorded in the other rootstocks, i.e.: 27, 21, 21, 30, 21, and 33 nmole Pi/mg DW for RG2, RG6, RG17, RX1, VLACH and WIP3, respectively. To investigate the extent to which rootstocks relied on the AM fungus for biomass production and Pi accumulation, the walnut overall behaviours in response to pre-inoculation with *R. irregularis* were compared as dependent on rootstock and Pi supply in terms of mycorrhizal dependency (MD%), mycorrhizal growth response (MGR%), and mycorrhizal phosphate response (MGP%). MD, MGR and shoot or root MGP% were positively correlated to each other under the lowest Pi fertilization regime tested (P/10), irrespective of the developmental stage. These results are consistent with the observation that in plant substrates with low available Pi, mycorrhizal growth responses (MGR%) and mycorrhizal dependency (MD%) of the plant are correlated, which may reflect a greater carbon supply provided by the host to the AM fungus [38]. The extent to which walnut rootstocks depend on the AM fungus for dry matter production varied between rootstocks and the Pi fertilization regime. Irrespective of root colonization parameters and developmental stages, mycorrhizal growth depression in the rootstock WIP3 after inoculation with *R. irregularis* was observed with increasing Pi availability in the substrate. These results also hold true for the rootstocks RG2, RG6, and RX1 after *post*-acclimatization. As reviewed by Johri et al. [39], inferior or negative MGRs at high Pi availabilities have often been attributed to the high carbon costs of the AM symbiosis that are not counterbalanced by a net gain in Pi. Consistently, the MPRs were always lower in these rootstock genotypes in the PTOT condition than in the P/10 treatment.

Plant dependence on mycorrhiza arises when uptake by roots alone of mineral nutrients at a given level of supply is not sufficient [40], with root nutrient acquisition ability being largely dependent on the root surface [41]. Within this line, RX1, which showed the lowest root surface (1 cm^2^) after acclimatization in the NM-PTOT treatment relative to the other rootstocks, further displayed the highest mycorrhizal dependency (53.5%) after *post*-acclimatization under the P/10 condition. It is noteworthy that a large absorption area of the root system is an inherent property of the plant or can be induced by P deficiency [42]. This was the case for the RG2, RG6, RG17, RX1, VLACH and WIP3 in which the root surfaces were higher in the P/10 than in the PTOT fertilization regime in both NM and AM plants after *post*-acclimatization. A significant converse trend was noticed after acclimatization and *post*-acclimatization for VX211, which displayed lower root surfaces in the P/10 than in the PTOT fertilization regime in both NM and AM plants. Owing to significantly increased leaf Pi levels in response to the AM fungus in the P/10 treatment whatever the developmental stage, these results suggest that *R. irregularis* provided greater effects in nutrient uptake than in the increase of root absorption area [38].

To further assess the impact of pre-inoculation with *R. irregularis* on rootstock robustness and biomass distribution, we coined the mycorrhizal quality response (MQR%), which besides shoot and root dry weight matters, also integrated morphological traits such as plant height, stem diameter, and dry weight matter partitioning [43]. MQR% ranged from negative to positive values depending on the rootstock studied and the Pi supply. MQR, MGR and MD% positively correlated to each other under the lowest Pi fertilization regime tested (P/10), irrespective of the developmental stage. These results underline that when grown under Pi limitation, the rootstocks VLACH, VX211, and WIP3 showed a greater DQI than under full Pi fertilization regime. Higher DQIs in response to AM symbiosis have been previously reported in plants such as *Melia azedarach* and *Cannabis sativa*, relative to non-inoculated controls [44]. Results showed that the higher mycorrhizal quality response recorded in the P/10 than the in the PTOT fertilization regime was mainly driven by a higher root DW for WIP3, and a lower shoot to root dry matter ratio for RG2, VLACH, VX211 in AM plants than in non-inoculated controls after acclimatization, indicating a larger investment in root DW production. By contrast, after *post*-acclimatization, the increased shoot DW recorded for VLACH, VX211 and WIP3 in AM plants than in non-inoculated controls rather explained the higher MQR% recorded in the P/10 than the in the PTOT fertilization regime. Taken together, these results support the hypothesis that an increased ability to develop young roots is fundamental during the first acclimatization stage of walnut rootstocks [6], which may be increased by mycorrhization. On the opposite, when micropropagated plantlets have an already developed root system after the acclimatization stage, they mainly rely on above ground expansion to develop their photosynthetic capacity in order to provide enough carbon to feed the AM fungus, which in return will bring back nutrients, instead of allocating carbon to root system development.

In conclusion, the main finding of this study relates to the ability of a commercially available AM fungus to decrease walnut rootstock dependency on Pi fertilization at both the acclimatization and *post*-acclimatization stages, together with improving adequate sapling development and quality, especially in the hybrid rootstocks VLACH, VX211, and WIP3. Our results also revealed the substantial variation of the response to mycorrhization between the different rootstock genotypes. Such variability in the response to AM symbiosis can be explored to improve the production of inoculated micropropagated rootstock in nurseries and to test their interest for agroforestry systems.

## Materials and Methods

### Biological material

Seven micropropagated rootstocks were used: *J. regia* cultivars RG2, RG6, RG17 and *Juglans* hybrids RX1 (*J. microcarpa x J. regia*), Vlach (*J. hindsi x J. regia*), VX211 (*J. hindsii x J. regia*), WIP3 ((*J. hindsii x J. regia) x J. regia)*). Propagation of the rootstocks was carried out on DKW medium [5] with 0.4 mg.L^−1^ 6-benzylaminopurine and 0.2 mg.L^−1^ indole-3-butiric acid. Rooting was initiated on MS medium modified by Jay-Allemand et al. [45] with 10 mg.L^−1^ Indole-3-Butiric acid for 7 days in the dark. After initiation, cuttings were transferred to modified MS medium without growth regulator for root expression for 4 weeks in the light before acclimatization (**Figure. S1A**). The AM fungus *R. irregularis* DAOM197198 (also known as DAOM181602) was purchased from Agronutrition (Labège, France) as an axenic spore suspension containing 200 spores.mL^−1^.

### Experiment 1. Nutrition, growth and quality of micropropagated walnut rootstocks after acclimatization

Rootstocks were placed for two months in individual alveolar plates in closed mini-greenhouses (Rapid Grow, Nortene; **Figure S1B**). Each alveolus was filled with an autoclaved (120 °C, 2 × 6h) silica sand (Biot): zeolite (Symbion) substrate (1:1, v : v). Two days after their transfer in the sand-zeolite substrate, plants were inoculated twice or not with 1 mL of the fungal spore suspension. Plants were sprayed with water twice a week to maintain humidity and fertilized once a week with 15 mL of either a low-Pi (P/10, 0.13 mM NaH_2_PO4, 2H_2_O) or a full-strength Pi (PTOT, 1.3 mM NaH_2_PO_4_, 2H_2_O) Hoagland solution. Walnut cultures were maintained for two months in a growth chamber (16 h photoperiod, 23 °C/18 °C day/night, 100% relative humidity, 220 μE m^−2^ s^−1^) before harvest (**Figure S1C**). The experiment was performed with a number of replicates per treatment ranging from height to 15.

### Experiment 2. Nutrition, growth and quality of micropropagated walnut rootstocks after *post*-acclimatization

Rootstocks were placed for *ex vitro* acclimatization in closed mini-greenhouses filled with the sand-zeolite substrate and grown in growth chamber conditions as described in experiment 1 (**Figure S1D**). They were sprayed twice a week with water to maintain humidity and fertilized once a week with 15 mL per plant of a NPK 6-4-6 solution containing 2% of magnesium and oligoelements (Algoflash). During two months, plantlets were gradually exposed to reduced relative humidity by progressively removing the box covers during a further month, then rootstocks were transplanted in individual pots containing 3 L of the sterilized the sand-zeolite substrate (**Figure S1E**). Two days after their transfer, plants were inoculated twice or not with 1 mL of the fungal spore suspension. Plants were fertilized once a week with 15 mL per plant of either a low-Pi or a full-strength Pi Hoagland solution during two months before harvest (**Figure S1F**). The experiment was performed with a number of replicates per treatment ranging from height to 15.

### Measurement of plant growth and nutritional parameters

All the parameters depicted in **Figure S2** were measured two months after inoculation. Root scans were analysed with ImageJ to assess root surface areas. To determine dry matter weights, shoot and root subsamples were weighed fresh, dried for two days at 80 °C and weighed. To measure Pi contents, dry root and shoot samples were finely ground using a Retsch Mixer ball mill (Haan, Germany). Free Pi was determined by measuring absorbance at 650 nm using the BIOMOL Green™ Reagent (Enzo Life Sciences, Villeurbanne, France).

### Quantification of AM root colonization

At harvest, root samples were randomly collected from each plant, dried for two days at 80°C, cleared at 90 °C for 30 min in 10% (w/v) KOH. After bleaching with three drops of 30% H_2_O_2_ [46], samples were stained in a 0.05% (w/v) trypan blue solution in lactic acid-glycerol-demineralised water (1:1:1) for 30 min at 90 °C. For each sample, thirty fragments (10 mm long, in glycerol) were observed and frequency of mycorrhization (F %), percentage of root cortex colonisation (M %), percentage of arbuscules (A %) were calculated according to [47].

### Metrics and statistical analyses

The Dickson Quality Index (DQI) was calculated individually for each plant according to the following formula [14]: DQI = (Shoot DW (g) + Root DW (g))/ (Plant height (cm)/Stem diameter (mm) + Shoot DW/Root DW). The mycorrhizal dependency (MD) was calculated based on the average of total dry matter content of the plants according to [48]: MD = 100 x (Dry matter of mycorrhizal plant – dry matter of non-mycorrhizal plant)/dry matter of mycorrhizal plant). The mycorrhizal growth response (MGR) was calculated based on the average total dry matter content of the plants according to [49]: MD = 100 x (Dry matter of mycorrhizal plant - dry matter of non-mycorrhizal plant)/dry matter of non-mycorrhizal plant). The mycorrhizal quality response (MQR) was calculated in the same way, with values of DQI in the place of dry matter in the latter equation. We determined the effect of pre-inoculation with *R. irregularis* on the shoot and root Pi content of the plant (mycorrhizal phosphate response; MPR) according to the following formula [50]: MPR = 100 x (Pi content of mycorrhizal plant - Pi content of non-mycorrhizal plant)/Pi content of non-mycorrhizal plant). Results were explored through a principal component analysis (PCA) with XLSTAT (Microsoft). As mycorrhizal colonization parameters represent percentages, all data were arcsine square root-transformed. Significant differences (*p* < 0.05) between transformed values recorded between the two contrasting Pi fertilization regimes were analysed using XLSTAT by the Welch-test (degrees of freedom = n-1), which is compatible with unequal variances between groups [51]. For each rootstock, the impacts of the pre-inoculation treatment and the Pi fertilization regime on the growth and nutrition parameters were analysed using the Kruskal-Wallis test. When the Kruskal-Wallis test was significant (*p* < 0.05), a *post*-hoc analysis was performed to determine the extent to which variables differ from each other by using the False Discovery Rate *p*-value adjustment method (FDR, *p*-adj < 0.05). The three latter tests were performed under Rstudio.

## Acknowledgements

Emma Mortier’s PhD thesis was funded by The French Ministry of Agriculture and Food. We would like to thank all the people who provided us with invaluable assistance in carrying out this work.

## Author contributions

E.M., F.M.-L., G.R., and O.L. planned, designed, perform the experiments and analysed data, S.J. analysed data, L.J. managed *in vitro* growth of walnut rootstocks, E.M., F.M.-L., G.R., and O.L. wrote the manuscript.

## Data availability

The data supporting the findings of this study are available from the corresponding authors upon reasonable request.

## Conflicts of interests

The authors declare no conflicts of interest.

## Supplementary information

**Supplemental Figure S1.**
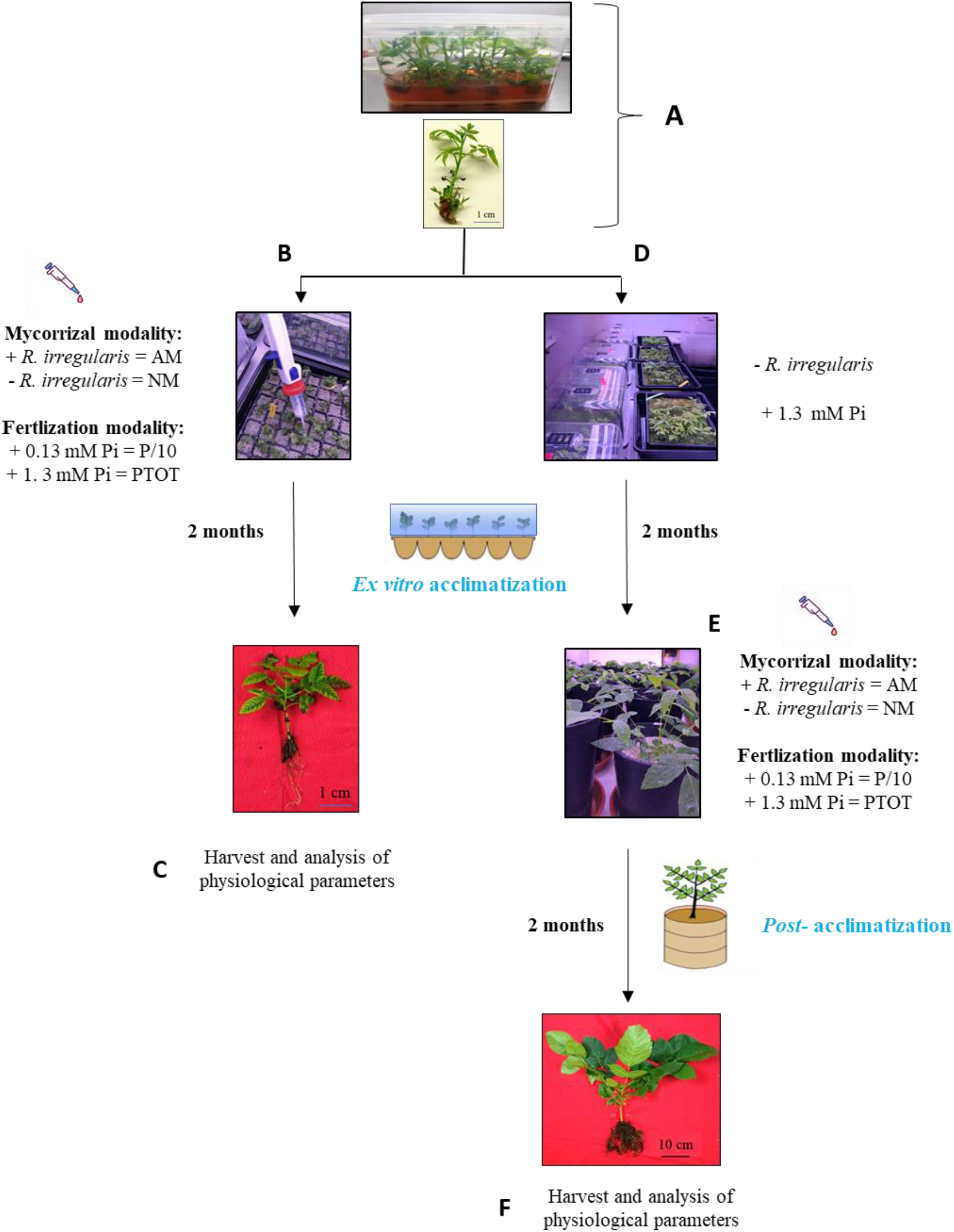
Experimental design. Micropropagated walnut rootstocks upon receipt and before their transfer into mini-greenhouses for acclimatization (A). For experiment 1 (B), plants were treated at the beginning of the acclimatization phase and physiological parameters were recorded two months after (C). For experiment 2 (D), plants were acclimatized during two months before their transfer in pots (E). Plants were treated at the beginning of the *post*-acclimatization phase and physiological parameters were recorded two months after (F).

**Supplementary Figure S2.**
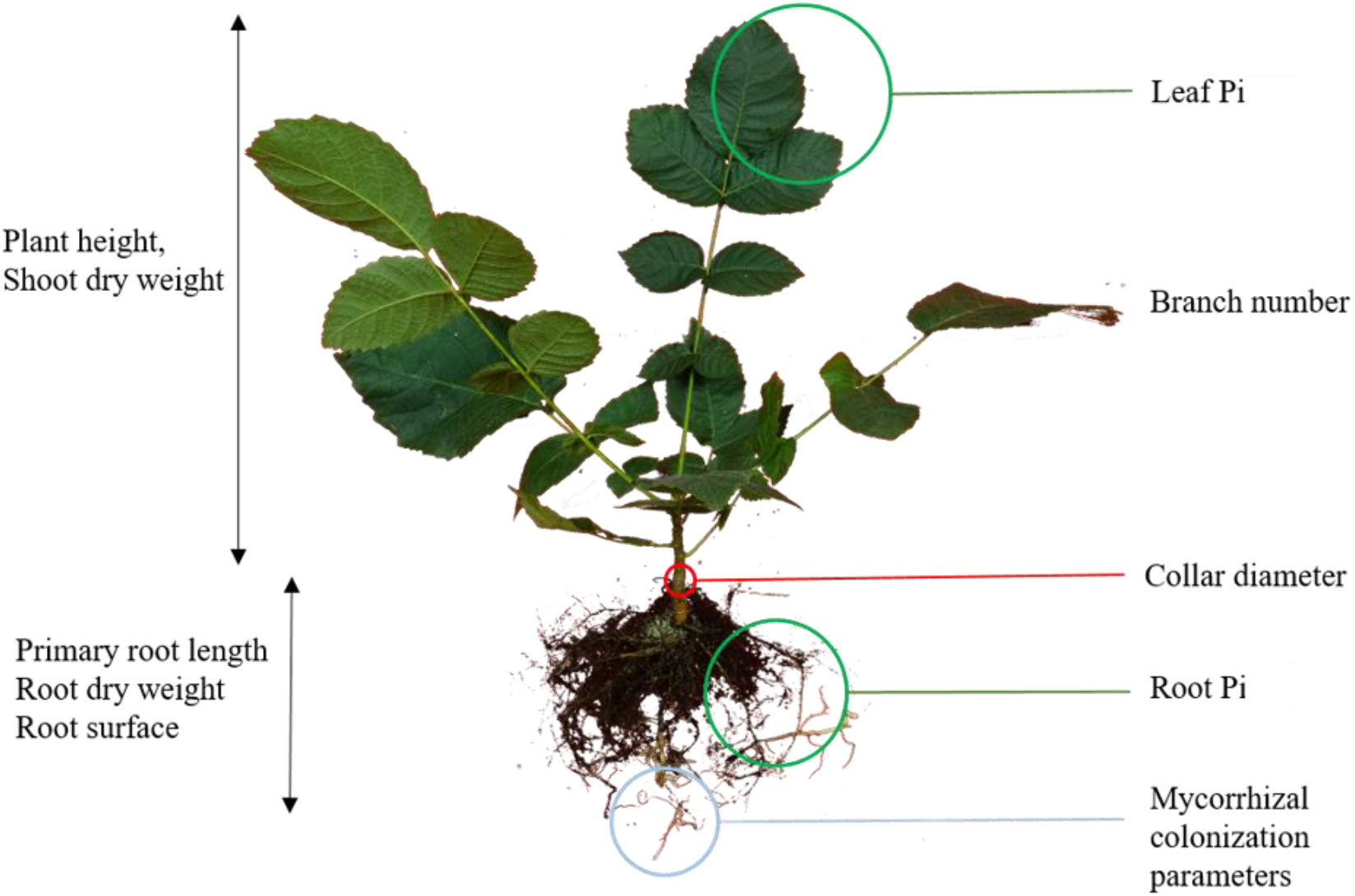
Nutritional and growth parameters analysed in this study.

**Supplementary data table S1.** Growth, quality and nutritional parameters of walnut rootstocks pre-inoculated with *R. irregularis* after two months of acclimatization under two different Pi supplies. Values correspond to the mean (±SE) of height to 15 replicates per treatment. Different letters indicate statistical differences between groups (*p* < 0.05) calculated as described in the material and methods section. AM: mycorrhizal; NM: non-mycorrhizal; DW: dry weight; DQI: Dickson Quality Index.

|  | P/10 |  | PTOT |  |  |
| --- | --- | --- | --- | --- | --- |
| **RG2** | AM | NM | AM | NM | *p* value |
| Root length (cm) | 8.1±0.7 | 8.0±0.7 | 8.0±0.9 | 10.7±1.7 | 0.37 |
| Collar (mm) | 2.5±0.2 | 2.2±0.2 | 2.9±0.2 | 2.7±0.2 | 0.15 |
| Height (cm) | 17.6±1.6 | 17.5±1.2 | 23.0±1.9 | 18.8±5.0 | 0.32 |
| Branch number | 18.0±1.2 | 16.6±2.0 | 20.6±1.2 | 13.3±3.8 | 0.23 |
| Root DW (g) | 0.07±0.01 | 0.05±7e<sup>-3</sup> | 0.08±0.02 | 0.07±0.02 | 0.63 |
| Shoot DW (g) | 0.04±5.6e<sup>-3</sup> b | 0.05±6.6e<sup>-3</sup> ab | 0.07±7.e<sup>-3</sup> a | 0.06±8.3e<sup>-3</sup> ab | **0.03** |
| Total DW(g) | 0.12±0.02 | 0.1±0.01 | 0.15±0.02 | 0.13±0.02 | 0.37 |
| Shoot/root DW ratio | 5.9±0.1 | 8.9±0.1 | 10.1±0.1 | 6.9±0.2 | 0.07 |
| DQI | 0.02±0.01 | 0.01±0.02 | 0.02±0.02 | 0.02±0.01 | 0.65 |
| Root surface (cm<sup>2</sup>) | 3.0±0.5 | 3.6±0.7 | 2.3±0.1 | 3.4±0.5 | 0.55 |
| Leaf Pi nmoles/mg DW | 27.2±7.0 bc | 16.2±2.81 c | 32.1±4.4 ab | 55.5 ±4.4 a | **0.02** |
| **RG6** | AM | NM | AM | NM | *p* value |
| Root length (cm) | 7.56±1.4 | 8.38±1.2 | 5.04±1.4 | 7.56±1.0 | 0.36 |
| Collar (mm) | 2.33±0.1 | 2.34±0.1 | 2.17±0.3 | 2.33±0.1 | 1.0 |
| Height (cm) | 6.94±1.3 | 8.98±0.8 | 7.98±1.4 | 10.96±1.0 | 0.08 |
| Branch number | 12.2±1.7 | 13.2±1.2 | 9.6±2.2 | 10.43±1.7 | 0.4 |
| Root DW (g) | 0.06±0.01 ab | 0.05±6.0e<sup>-3</sup> ab | 0.08±0.02 a | 0.03±6.4e<sup>-3</sup> b | **0.04** |
| Shoot DW (g) | 0.2±0.04 | 0.2±0.07 | 0.2±0.07 | 0.2±0.03 | 0.97 |
| Total (g) | 0.2±0.05 | 0.3±0.07 | 0.2±0.08 | 0.2±0.03 | 0.95 |
| Shoot/root DW ratio | 2.9±0.6 | 4.3±1.0 | 2.6±0.7 | 5.6±0.7 | 0.07 |
| DQI | 0.04±0.004 a | 0.03±0.004 ab | 0.04±0.001 a | 0.02±0.003 b | **0.03** |
| Root surface (cm<sup>2</sup>) | 3.1±0.5 | 1.9±0.4 | 1.5±0.6 | 2.1±0.4 | 0.4 |
| Leaf Pi nmoles/mg DW | 36.9±7.2 | 40.8±9.4 | 48.6±5.1 | 49.1±6.7 | 0.47 |
| **RG17** | AM | NM | AM | NM | *p* value |
| Root length (cm) | 9.2±1.4 | 9.9±0.6 | 9.2±1.1 | 10.6±0.6 | 0.94 |
| Collar (mm) | 2.2±0.1 | 2.5±0.1 | 2.42±0.2 | 2.1±0.2 | 0.15 |
| Height (cm) | 11.9±1.2 | 10.7±1.1 | 14.4±2.2 | 13.3±1.6 | 0.61 |
| Branch number | 19.4±1.8 | 16.9±0.9 | 20.2±1.7 | 18.9±1.4 | 0.43 |
| Root DW (g) | 0.12±0.03 <sup>a</sup> | 0.06±0.01 <sup>b</sup> | 0.15±0.03 <sup>a</sup> | 0.08±0.02 <sup>b</sup> | **0.02** |
| Shoot DW (g) | 0.4±0.05 | 0.3±0.05 | 0.5±0.1 | 0.3±0.02 | 0.44 |
| Total DW (g) | 0.5±0.06 | 0.4±0.06 | 0.6±0.13 | 0.5±0.2 | 0.30 |
| Shoot/root DW ratio | 3.8±0.5 <sup>b</sup> | 6.7±1.0 <sup>a</sup> | 3.7±0.7 <sup>b</sup> | 6.6±1.1 <sup>a</sup> | **0.02** |
| DQI | 0.06±0.001 <sup>ab</sup> | 0.04±0.007 <sup>bc</sup> | 0.07±0.01 <sup>a</sup> | 0.04±0.02 <sup>c</sup> | **0.01** |
| Root surface (cm<sup>2</sup>) | 4.4±0.6 | 3.8±0.8 | 1.8±1.0 | 3.39±1.2 | 0.42 |
| Leaf Pi nmoles/mg DW | 36.5±5.6 | 36.5±6.1 | 56.5±8.9 | 38.35±4.6 | 0.23 |
| **RX1** | AM | NM | AM | NM | *p* value |
| Root length (cm) | 9.2±0.7 <sup>ab</sup> | 7.9±0.6 <sup>b</sup> | 9.6±0.5 <sup>ab</sup> | 10.4±0.5 <sup>a</sup> | **0.03** |
| Collar (mm) | 2.7±0.1 | 2.6±0.1 | 2.5±0.1 | 2.5±0.1 | 0.3 |
| Height (cm) | 12.44±0.4 | 12.5±0.7 | 13.7±0.8 | 11.7±1.0 | 0.3 |
| Branch number | 12.4±0.6 | 11.4±0.5 | 12.2±0.4 | 11.0±0.6 | 0.27 |
| Root DW (g) | 0.08±7.4e<sup>-3</sup> | 0.1±0.03 | 0.05±3.6e<sup>-3</sup> | 0.05±5.5e<sup>-3</sup> | 0.5 |
| Shoot DW (g) | 0.3±0.03 | 0.2±0.03 | 0.2±0.02 | 0.2±0.03 | 0.32 |
| Total DW (g) | 0.3±0.03 | 0.3±0.05 | 0.3±0.02 | 0.3±0.03 | 0.7 |
| Shoot/root DW ratio | 3.8±0.4 <sup>b</sup> | 6.6±0.3 <sup>a</sup> | 3.7±0.3 <sup>b</sup> | 6.6±0.3 <sup>a</sup> | **0.01** |
| DQI | 0.04±0.003 | 0.04±0.01 | 0.03±0.002 | 0.03±0.02 | 0.1 |
| Root surface (cm<sup>2</sup>) | 1.8±0.2 | 1.4±0.2 | 1.1±0.1 | 1.0±0.1 | 0.1 |
| Leaf Pi nmoles/mg DW | 20.5±1.6 <sup>b</sup> | 16.2±2.8 <sup>b</sup> | 32.1±3.0 <sup>b</sup> | 35.4±2.8 <sup>a</sup> | **<0.01** |
| **VLACH** | AM | NM | AM | NM | *p* value |
| Root length (cm) | 7.5±0.3 <sup>bc</sup> | 6.0±0.4 <sup>c</sup> | 9.0 ±0.7 <sup>b</sup> | 11.5±0.8 <sup>a</sup> | **<0.01** |
| Collar (mm) | 3.3±0.2 <sup>a</sup> | 2.7±0.2 <sup>b</sup> | 2.6±0.1 <sup>b</sup> | 2.7±0.1 <sup>b</sup> | **0.02** |
| Height (cm) | 9.1±0.6 <sup>ab</sup> | 6.7±0.4 <sup>b</sup> | 10.8±0.7 <sup>a</sup> | 9.7±0.9 <sup>a</sup> | **<0.01** |
| Branch number | 10.7±0.5 <sup>ab</sup> | 9.±30.5 <sup>b</sup> | 12.1±0.5 <sup>a</sup> | 11.5±0.9 <sup>ab</sup> | **0.04** |
| Root DW (g) | 0.1±9.1e<sup>-3</sup> <sup>a</sup> | 0.05±6.3 e<sup>-3</sup> <sup>c</sup> | 0.08±7.9e<sup>-3</sup> <sup>b</sup> | 0.1±0.01 <sup>a</sup> | **<0.01** |
| Shoot DW (g) | 0.3±0.04 <sup>a</sup> | 0.2±0.03 <sup>b</sup> | 0.3±0.04 <sup>a</sup> | 0.4±0.06 <sup>a</sup> | **<0.01** |
| Total DW (g) | 0.4±0.05 <sup>a</sup> | 0.2±0.03 <sup>b</sup> | 0.4±0.04 <sup>a</sup> | 0.5±0.06 <sup>a</sup> | **<0.01** |
| Shoot/root DW ratio | 2.5±0.4 <sup>b</sup> | 3.7±0.4 <sup>ab</sup> | 4.4±0.3 <sup>a</sup> | 4.2±0.3 <sup>a</sup> | **<0.01** |
| DQI | 0.07±0.005 <sup>a</sup> | 0.04±0.005 <sup>b</sup> | 0.04±0.004 <sup>b</sup> | 0.06±0.006 <sup>ab</sup> | **<0.01** |
| Root surface (cm<sup>2</sup>) | 1.7±0.1 | 1.2±0.2 | 1.4±0.1 | 1.4±0.1 | 0.15 |
| Leaf Pi nmoles/mg DW | 81.7±11.4 <sup>a</sup> | 68.3±3.3 <sup>b</sup> | 81.3±4.7 <sup>a</sup> | 66.2±3.3 <sup>b</sup> | **<0.01** |
| **VX211** | AM | NM | AM | NM | *p* value |
| Root length (cm) | 5.6±0.3<sup>b</sup> | 6.0±0.9<sup>b</sup> | 12.3±1.6<sup>a</sup> | 12.2±1.2<sup>a</sup> | **<0.01** |
| Collar (mm) | 2.1±0.1<sup>ab</sup> | 2.2±0.1<sup>b</sup> | 3.0±0.2<sup>a</sup> | 3.1±0.2<sup>a</sup> | **<0.01** |
| Height (cm) | 7.8±0.4 | 7.5±0.6 | 7.0±0.9 | 7.0±0.8 | 0.2 |
| Branch number | 12.0±0.5 | 10.3±0.6 | 10.9±1.1 | 10.1±1.3 | 0.2 |
| Root DW (g) | 0.09±8.8e<sup>-3</sup><sup>a</sup> | 0.05±3. e<sup>-3</sup><sup>b</sup> | 0.1±0.03<sup>a</sup> | 0.2±0.03<sup>a</sup> | **<0.01** |
| Shoot DW (g) | 0.1±0.01<sup>a</sup> | 0.1±0.02<sup>a</sup> | 0.05±9.5e<sup>-3</sup><sup>b</sup> | 0.06±7.8e<sup>-3</sup><sup>b</sup> | **<0.01** |
| Total DW (g) | 0.2±0.02 | 0.2±0.02 | 0.2±0.04 | 0.2±0.04 | 0.3 |
| Shoot/root DW ratio | 1.4±0.1<sup>b</sup> | 2.6±0.4<sup>a</sup> | 0.6±0.1<sup>c</sup> | 0.5±0.1<sup>c</sup> | **<0.01** |
| DQI | 0.05±0.004<sup>b</sup> | 0.03±0.002<sup>b</sup> | 0.08±0.02<sup>ab</sup> | 0.1±0.002<sup>a</sup> | **0.01** |
| Root surface (cm<sup>2</sup>) | 1.1±0.1<sup>b</sup> | 0.9±0.1<sup>b</sup> | 2.8±0.3<sup>a</sup> | 6.5±0.9<sup>a</sup> | **<0.01** |
| Leaf Pi nmoles/mg DW | 43.8±3.9<sup>a</sup> | 29.7±3.8<sup>b</sup> | 42.4±3.2<sup>a</sup> | 26.7±4.5<sup>b</sup> | **<0.01** |
| **WIP3** | AM | NM | AM | NM | *p* value |
| Root length (cm) | 7.0±0.4 | 7.7±0.6 | 9.0±0.8 | 9.8±0.9 | 0.07 |
| Collar (mm) | 2.8±0.1 | 2.7±0.1 | 2.9±0.2 | 3.1±0.2 | 0.26 |
| Height (cm) | 8.34±0.5<sup>bc</sup> | 7.8±0.6<sup>c</sup> | 12.3±1.0<sup>a</sup> | 11.5±1.0<sup>ab</sup> | **<0.01** |
| Branch number | 10.6±0.4<sup>b</sup> | 9.7±0.6<sup>b</sup> | 13.1±0.7<sup>a</sup> | 14.0±1.0<sup>a</sup> | **<0.01** |
| Root DW (g) | 0.09±5.8e<sup>-3</sup> | 0.06±7.6e<sup>-3</sup> | 0.07±0.01 | 0.09±0.02 | 0.08 |
| Shoot DW (g) | 0.5±0.2 | 0.2±0.04 | 0.2±0.05 | 0.3±0.07 | 0.55 |
| Total DW (g) | 0.5±0.2 | 0.3±0.05 | 0.3±0.07 | 0.4±0.08 | 0.7 |
| Shoot/root DW ratio | 5.1±2.4 | 3.5±0.6 | 3.7±0.3 | 3.5±0.3 | 0.56 |
| DQI | 0.06±0.004 | 0.04±0.006 | 0.04±0.009 | 0.05±0.01 | 0.65 |
| Root surface (cm<sup>2</sup>) | 1.6±0.2<sup>b</sup> | 2.0±0.3<sup>b</sup> | 3.6±0.4<sup>a</sup> | 4.6±0.5<sup>a</sup> | **<0.01** |
| Leaf Pi nmoles/mg DW | 69.3±4.0<sup>a</sup> | 64±5.1<sup>b</sup> | 67.9±3.8<sup>a</sup> | 53.0±4.7<sup>b</sup> | **<0.01** |

**Supplementary data table S2.** Mycorrhizal rootstock responses to pre-inoculation with *R. irregularis* after two months of acclimatization under two different Pi supplies. Values calculated from to the mean of height to 15 replicates per mycorrhizal and non-mycorrhizal plants for each Pi supply, were categorized as follows: −5 %< neutral< 5% (grey values), negative :< −5% (blue values), 5 %< positive< 50% (pink values); highly positive? 50% (red values). MD%, mycorrhizal dependency; MGR%, mycorrhizal growth response, MPRs%, mycorrhizal shoot phosphate response, MQR%, mycorrhizal quality response.

|  | <u>RG2</u> |  | <u>RG6</u> |  | <u>RG17</u> |  | <u>RX1</u> |  | <u>VLACH</u> |  | <u>VX211</u> |  | <u>WIP3</u> |  |
| --- | --- | --- | --- | --- | --- | --- | --- | --- | --- | --- | --- | --- | --- | --- |
|  | P/10 | PTOT | P/10 | PTOT | P/10 | PTOT | P/10 | PTOT | P/10 | PTOT | P/10 | PTOT | P/10 | PTOT |
| MGR% | 17 | 14 | -12 | 30 | 26 | 25 | 4 | 4 | 74 | -15 | 27 | -18 | 113 | -20 |
| MD% | 15 | 12 | -14 | 23 | 21 | 20 | 4 | 4 | 42 | -17 | 21 | -22 | 53 | -25 |
| MQR% | 100 | 0 | 33 | 100 | 50 | 75 | 0 | 0 | 75 | -33 | 67 | -20 | 50 | -20 |
| MPRs% | 34 | -32 | -10 | -1 | 33 | 146 | 90 | -9 | 80 | -8 | 47 | 59 | 169 | -36 |

**Supplementary data table S3.** Growth, quality and nutritional parameters of walnut rootstocks pre-inoculated with *R. irregularis* after two months of *post*-acclimatization under two different Pi supplies. Values correspond to the mean (±SE) of height to 15 replicates per treatment. Different letters indicate statistical differences between groups (*p* < 0.05) calculated as described in the material and methods section. AM: mycorrhizal; NM: non-mycorrhizal; DW: dry weight; DQI: Dickson Quality Index.

|  | P/10 |  | PTOT |  |  |
| --- | --- | --- | --- | --- | --- |
| **RG2** | AM | NM | AM | NM | *p* value |
| Root length (cm) | 20.2±1.1 <sup>c</sup> | 31.4±2.3 <sup>ab</sup> | 28.1±2.3 <sup>b</sup> | 42.7±5.9 <sup>a</sup> | **<0.01** |
| Collar (mm) | 6.9±0.2 <sup>c</sup> | 7.1±0.4 <sup>bc</sup> | 8.3±0.5 <sup>ab</sup> | 9.2±0.8 <sup>a</sup> | **0.02** |
| Height (cm) | 26.3±1.3a | 19.0±2.0 | 27.8±3.4 | 24.4±5.4 | 0.14 |
| Branch number | 9.9±0.3 | 9.9±0.2 | 9.6±0.4 | 9.3±0.4 | 0.92 |
| Root DW (g) | 1.9±0.1 <sup>b</sup> | 2.1±0.3 <sup>b</sup> | 5.22±1.1 <sup>a</sup> | 10.0±3.3 <sup>a</sup> | **<0.01** |
| Shoot DW (g) | 4.5±1.8 | 4.1±0.9 | 7.3±1.9 | 6.6±1.9 | 0.18 |
| Total DW(g) | 5.8±1.7 <sup>b</sup> | 6.2±1.0 <sup>b</sup> | 12.5±2.7 <sup>a</sup> | 16.6±4.8 <sup>a</sup> | **<0.01** |
| Shoot/root DW ratio | 2.8±1.4 <sup>ab</sup> | 2.2±0.4 <sup>a</sup> | 1.4±0.3 <sup>ab</sup> | 0.84±0.2 <sup>b</sup> | 0.05 |
| DQI | 0.8±0.1 <sup>b</sup> | 1.2±0.3 <sup>b</sup> | 2.6±0.5 <sup>a</sup> | 6.6±2.6 <sup>a</sup> | **<0.01** |
| Root surface (cm<sup>2</sup>) | 692.2±70.2 <sup>a</sup> | 593.8±35.5 <sup>ab</sup> | 433.3±68.7 <sup>bc</sup> | 314.0±47.1 <sup>c</sup> | **0.01** |
| Root Pi nmoles<br>/mg DW | 17.7±1.2 | 14.54±1.5 | 27.5±5.76 | 19.72±4.09 | 0.09 |
| Leaf Pi nmoles<br>/mg DW | 36.7±2.0 <sup>a</sup> | 22.7±2.7 <sup>b</sup> | 37.8±7.2 <sup>a</sup> | 22.2±3.2 <sup>b</sup> | **0.02** |
| **RG6** | AM | NM | AM | NM | *p* value |
| Root length (cm) | 27.1±1.5 | 29.6±1.6 | 33.9±3.7 | 31.6±2.0 | 0.64 |
| Collar (mm) | 7.5±0.5 | 8.0±0.4 | 7.5±0.8 | 7.8±0.6 | 0.69 |
| Height (cm) | 17.1±2.4 | 14.9±3.0 | 13.4±2.8 | 12.6±2.0 | 0.69 |
| Branch number | 8.7±1.0 | 9.3±0.8 | 7.2±0.5 | 6.7±0.4 | 0.11 |
| Root dw (g) | 3.0±0.6 | 2.35±0.3 | 6.4±1.4 | 7.4±1.8 | 0.09 |
| Shoot dw (g) | 5.3±0.7 | 5.1±1.1 | 5.2±1.5 | 4.3±1.0 | 0.65 |
| Total dw (g) | 8.3±1.0 | 7.5±1.0 | 11.6±2.5 | 11.6±2.7 | 0.97 |
| Shoot/root DW ratio | 2.5±0.5 <sup>a</sup> | 2.7±0.7 <sup>a</sup> | 1.0±0.3 <sup>b</sup> | 0.7±0.1 <sup>b</sup> | **<0.01** |
| DQI | 2.2±0.4 | 2.2±0.3 | 6.2±1.9 | 6.5±1.7 | 0.27 |
| Root surface (cm<sup>2</sup>) | 510.6±77.8 | 629.5±67.2 | 364.7±51.8 | 454.5±46.5 | 0.65 |
| Root Pi nmoles<br>/mg DW | 19±1.7 | 16.7±1.3 | 20.8±3.0 | 21.4±3.0 | 0.87 |
| Leaf Pi nmoles<br>/mg DW | 27.6±2.9 | 26.1±3.0 | 27.9±5.0 | 40.0±7.0 | 0.31 |
| **RG17** | AM | NM | AM | NM | *p* value |
| Root length (cm) | 31.6±2.4 | 42.3±2.3 | 39.7±4.3 | 47.3±6.0 | 0.15 |
| Collar (mm) | 6.4±0.3 | 6.5±0.4 | 7.3±0.8 | 6.8±0.7 | 0.98 |
| Height (cm) | 13.5±1.9 <sup>a</sup> | 8.0±1.3 <sup>b</sup> | 15.6±1.4 <sup>a</sup> | 11.8±1.4 <sup>ab</sup> | **<0.01** |
| Branch number | 11.6±0.9 <sup>a</sup> | 9.7±0.7 <sup>ab</sup> | 8.1±0.7 <sup>b</sup> | 8.4±1.0 <sup>b</sup> | **0.02** |
| Root DW (g) | 3.0±0.4 | 3.6±0.6 | 5.7±1.4 | 5.3±1.2 | 0.62 |
| Shoot DW (g) | 5.3±0.7 | 3.7±0.9 | 5.6±2.11 | 3.6±1.1 | 0.24 |
| Total DW (g) | 8.3±0.8 | 7.3±1.4 | 11.4±3.2 | 9.0±2.0 | 0.7 |
| Shoot/root DW ratio | 2.03±0.3 <sup>a</sup> | 1.0±0.1 <sup>ab</sup> | 0.8±0.2 <sup>b</sup> | 0.74±0.1 <sup>b</sup> | **<0.01** |
| DQI | 2.35±0.3 | 3.2±0.4 | 3.9±1.1 | 4.46±1.3 | 0.6 |
| Root surface (cm<sup>2</sup>) | 629.2±52.2 | 804.3±32.7 | 445.4±46.7 | 590.9±71.0 | 0.5 |
| Root Pi nmoles<br>/mg DW | 18.6±1.5 <sup>b</sup> | 13.2±0.9 <sup>c</sup> | 30.2±1.0 <sup>a</sup> | 27.0±4.4 <sup>b</sup> | **<0.01** |
| Leaf Pi nmoles<br>/mg DW | 35.5±2.1 <sup>b</sup> | 27.1±3.7 <sup>b</sup> | 102.7±8.9 <sup>a</sup> | 98.5±9.5 <sup>a</sup> | **<0.01** |
| **RX1** | AM | NM | AM | NM | *p* value |
| Root length (cm) | 38.6±2.1 | 40.5±3.4 | 34.87±1.6 | 35.3±1.9 | 0.07 |
| Collar (mm) | 5.5±0.4 <sup>a</sup> | 4.7±0.4 <sup>ab</sup> | 4.9±0.2 <sup>ab</sup> | 3.9±0.2 <sup>b</sup> | **0.02** |
| Height (cm) | 18.1±1.8 <sup>a</sup> | 16.6±1.1 <sup>ab</sup> | 13.2±1.1 <sup>b</sup> | 14.9±1.2 <sup>ab</sup> | 0.05 |
| Branch number | 6.1±0.6 | 6.8±0.7 | 5.6±0.2 | 5.8±0.4 | 0.53 |
| Root DW (g) | 1.7±0.4 | 1.6±0.4 | 1.4±0.3 | 2.0±0.5 | 0.3 |
| Shoot DW (g) | 2.7±0.6 <sup>a</sup> | 1.8±0.5 <sup>a</sup> | 1.8±0.4 <sup>a</sup> | 1.0±0.2 <sup>b</sup> | **<0.01** |
| Total DW (g) | 4.3±1.0 <sup>a</sup> | 2.0±0.6 <sup>b</sup> | 3.2±0.8 <sup>a</sup> | 3.0±0.2 <sup>a</sup> | **<0.01** |
| Shoot/root DW ratio | 2.5±0.6 <sup>a</sup> | 1.4±0.4 <sup>a</sup> | 1.9±0.5 <sup>a</sup> | 0.6±0.1 <sup>b</sup> | **<0.01** |
| DQI | 0.8±0.2 <sup>a</sup> | 0.3±0.1 <sup>b</sup> | 0.8±0.2 <sup>a</sup> | 0.2±0.1 <sup>b</sup> | **<0.01** |
| Root surface (cm<sup>2</sup>) | 501.4±67.8 <sup>a</sup> | 532.2±66.3 <sup>a</sup> | 149.2±40.3 <sup>b</sup> | 98.3±9.5 <sup>b</sup> | **<0.01** |
| Root Pi nmoles<br>/mg DW | 17.6±2.1 | 18.3±3.1 | 21.2±1.4 | 16.93±2.1 | 0.24 |
| Leaf Pi nmoles<br>/mg DW | 26.7±2.3 | 23.8±2.0 | 36.5±4.03 | 32.2±6.0 | 0.14 |
| **VLACH** | AM | NM | AM | NM | *p* value |
| Root length (cm) | 29.4±2.2<sup>ab</sup> | 37.1±2.8<sup>a</sup> | 23.3±1.8<sup>b</sup> | 30.5±2.5<sup>ab</sup> | **<0.01** |
| Collar (mm) | 7.2±0.7 | 6.4±0.4 | 5.7±0.5 | 5.6±0.5 | 0.13 |
| Height (cm) | 12.2±1.4 | 9.5±0.7 | 11.1±2.0 | 10.8±1.8 | 0.53 |
| Branch number | 6.1±0.6 | 6.1±0.5 | 6.4±1.1 | 5.6±0.8 | 0.9 |
| Root DW (g) | 2.5±0.5<sup>ab</sup> | 3.0±0.6<sup>a</sup> | 1.0±0.2<sup>c</sup> | 1.5±0.4<sup>bc</sup> | **<0.01** |
| Shoot DW (g) | 3.4±0.7 | 2.23±0.5 | 2.4±0.7 | 1.99±0.6 | 0.35 |
| Total DW (g) | 5.9±1.0 | 5.2±1.0 | 3.4±0.8 | 3.5±0.9 | 0.1 |
| Shoot/root DW ratio | 1.8±0.2<sup>a</sup> | 0.7±0.1<sup>b</sup> | 1.1±0.5<sup>ab</sup> | 1.2±0.4<sup>ab</sup> | **<0.01** |
| DQI | 2.1±0.5<sup>a</sup> | 2.5±0.6<sup>a</sup> | 0.8±0.1<sup>b</sup> | 1.2±0.3<sup>ab</sup> | **0.02** |
| Root surface (cm<sup>2</sup>) | 122.8±18.2 | 108.5±14.6 | 63.9±12.7 | 82.2±10.1 | 0.09 |
| Root Pi nmoles | 28.6±4.4<sup>a</sup> | 15.9±1.8<sup>b</sup> | 33.0±5.6<sup>a</sup> | 15.9±2.1<sup>b</sup> | **0.02** |
| /mg DW |  |  |  |  |  |
| Leaf Pi nmoles | 36.5±5.4<sup>a</sup> | 14.9±1.9<sup>b</sup> | 33.0±6.1<sup>a</sup> | 47.0±5.8<sup>a</sup> | **<0.01** |
| /mg DW |  |  |  |  |  |
| **VX211** | AM | NM | AM | NM | *p* value |
| Root length (cm) | 45.9±2.3 | 48.7±2.1 | 45.5±3.1 | 44.5±4.6 | 0.32 |
| Collar (mm) | 6.8±0.7 | 6.5±0.6 | 6.0±0.4 | 6.2±0.5 | 0.98 |
| Height (cm) | 10.6±1.8 | 10.6±1.6 | 9.4±0.8 | 9.6±0.8 | 0.92 |
| Branch number | 6.6±0.6 | 5.4±0.7 | 5.4±0.4 | 5.4±0.5 | 0.64 |
| Root DW (g) | 6.7±1.3 | 5.2±1.2 | 8.9±1.7 | 9.1±2.2 | 0.33 |
| Shoot DW (g) | 3.7±0.8 | 3.1±0.7 | 2.2±0.7 | 2.5±0.6 | 1.0 |
| Total DW (g) | 9.6±2.0 | 8.0±1.7 | 11.1±2.0 | 11.6±2.8 | 0.6 |
| Shoot/root DW ratio | 0.5±0.06<sup>a</sup> | 0.5±0.05<sup>a</sup> | 0.3±0.03<sup>b</sup> | 0.3±0.1<sup>b</sup> | **<0.01** |
| DQI | 4.5±0.8 | 3.7±0.6 | 7.7±1.8 | 7.4±1.8 | 0.44 |
| Root surface (cm<sup>2</sup>) | 169.5±19.9<sup>b</sup> | 197.93±20.5<sup>b</sup> | 262.1±14.9<sup>a</sup> | 255.7±12.9<sup>a</sup> | **<0.01** |
| Root Pi nmoles | 18.0±1.0<sup>b</sup> | 10.5±1.1<sup>c</sup> | 45.9±6.4<sup>a</sup> | 36.2±4.4<sup>a</sup> | **<0.01** |
| /mg DW |  |  |  |  |  |
| Leaf Pi nmoles | 22.7±3.1<sup>b</sup> | 10.4±1.4<sup>c</sup> | 87.8±10.0<sup>a</sup> | 56.3±9.4<sup>a</sup> | **<0.01** |
| /mg DW |  |  |  |  |  |
| WIP3 | AM | NM | AM | NM | *p* value |
| Root length (cm) | 24.2±1.5 <sup>b</sup> | 25.6±1.4 <sup>b</sup> | 29.5±2.1 <sup>ab</sup> | 35.5±1.9 <sup>a</sup> | **<0.01** |
| Collar (mm) | 6.6±0.2 | 6.1±0.2 | 6.2±0.5 | 5.6±0.2 | 0.11 |
| Height (cm) | 32.2±2.6 <sup>a</sup> | 14.8±1.1 <sup>b</sup> | 31.0±4.5 <sup>a</sup> | 14.7±1.3 <sup>b</sup> | **<0.01** |
| Branch number | 7.5±0.4 <sup>a</sup> | 6.3±0.4 <sup>bc</sup> | 7.3±0.5 <sup>ab</sup> | 5.1±0.4 <sup>c</sup> | **<0.01** |
| Root DW (g) | 3.6±0.7 | 3.2±0.4 | 1.6±0.3 | 2.4±0.9 | 0.06 |
| Shoot DW (g) | 4.8±0.5 | 3.2±0.6 | 5.0±0.8 | 5.0±0.9 | **<0.01** |
| Total DW (g) | 8.8±1.2 <sup>a</sup> | 6.2±1.0 <sup>ab</sup> | 2.9±0.5 <sup>c</sup> | 4.0±1.5 <sup>bc</sup> | **<0.01** |
| Shoot/root DW ratio | 1.8±0.3 | 1.1±0.2 | 1.2±0.2 | 1.5±0.2 | 0.16 |
| DQI | 1.8±0.4 | 1.1±0.19 | 1.31±0.27 | 2.1±0.7 | 0.4 |
| Root surface (cm²) | 529.5±50.0 <sup>b</sup> | 651.4±41.2 <sup>a</sup> | 197.3±14.1 <sup>c</sup> | 146.1±15.2 <sup>c</sup> | **<0.01** |
| Root Pi nmoles | 17.5±1.4 <sup>b</sup> | 12.0±1.1 <sup>c</sup> | 31.3±4.3 <sup>a</sup> | 26.3±2.2 <sup>a</sup> | **<0.01** |
| /mg DW |  |  |  |  |  |
| Leaf Pi nmoles | 33.1±1.9 <sup>a</sup> | 32.4±1.6 <sup>a</sup> | 53.8±8.72 <sup>b</sup> | 52.7±7.8 <sup>b</sup> | 0.06 |
| /mg DW |  |  |  |  |  |

**Supplementary data table S4.** Mycorrhizal rootstock responses to pre-inoculation with *R. irregularis* after two months of *post*-acclimatization under two different Pi supplies. Values calculated from to the mean of height to 15 replicates per mycorrhizal and non-mycorrhizal plants for each Pi supply, were categorized as follows: −5 %< neutral< 5% (grey values), negative :< −5% (blue values), 5 %< positive< 50% (pink values); highly positive> 50% (red values). MD%, mycorrhizal dependency; MGR%, mycorrhizal growth response, MPRs%, mycorrhizal shoot phosphate response, MQR%, mycorrhizal quality response.

|  | RG2 |  | RG6 |  | RG17 |  | RX1 |  | VLACH |  | VX211 |  | WIP3 |  |
| --- | --- | --- | --- | --- | --- | --- | --- | --- | --- | --- | --- | --- | --- | --- |
|  | P/10 | PTOT | P/10 | PTOT | P/10 | PTOT | P/10 | PTOT | P/10 | PTOT | P/10 | PTOT | P/10 | PTOT |
| MGR% | -6 | -24 | 11 | 0 | 14 | 8 | 115 | 7 | 13 | -3 | 20 | -4 | 44 | -27 |
| MD% | -7 | -33 | 10 | 0 | 12 | 22 | 53 | 6 | 12 | -3 | 17 | -5 | 29 | -38 |
| MQR% | -33 | -61 | 0 | -5 | -27 | -13 | 166 | 300 | -16 | -33 | 22 | 4 | 64 | -38 |
| MPRs% | 77 | 88 | 10 | -15 | 85 | 62 | 68 | 104 | 257 | -15 | 161 | 37 | 53 | -2 |
| MPRr% | 10 | -27 | 46 | -16 | 109 | 14 | 2 | -123 | 50 | 38 | 121 | 41 | 64 | -21 |

## Notes

### Competing Interest Statement

The authors have declared no competing interest.

